# Phylogenomics, biogeography, and evolution in the American palm genus *Brahea*

**DOI:** 10.1101/467779

**Authors:** Craig F. Barrett, Brandon T. Sinn, Loren T. King, Jesus C. Medina, Christine D. Bacon, Sean C. Lahmeyer, Donald R. Hodel

## Abstract

**Background and Aims:** Slow rates of molecular evolution at low taxonomic levels hamper studies of relationships among species, and subsequent biogeographic and evolutionary analyses. An example is the genus *Brahea*, which is among the most poorly understood lineages of American palms and is characterized by a wide variety of growth forms and intermediate morphological features.

**Methods:** We generated approximately 400 kb of genome-scale data from all three genomes for the 11 currently described species of *Brahea* to infer phylogenetic relationships, reconstruct ancestral growth form, estimate ancestral geographic ranges, and test for niche equivalency among closely related species with geographic overlap.

**Key Results:** Relationships receive strong support, and conform to previous subgeneric assignments, except for placement of the dwarf species *B. moorei* within subgenus *Erythea.* Our robust phylogenetic hypothesis reveals trends in growth form including an overall increase in height in the *B. armata* clade, and independent evolution of dwarf forms from taller ancestors in the *B. pimo* and *B. dulcis* clades. Ancestral range estimation reveals roles of dispersal (e.g. *B. edulis* on Guadalupe Island) and sympatric speciation in some cases (e.g. in the *B. armata* clade), but is equivocal in others (e.g. in the *B. pimo clade*). We find evidence of niche non-equivalency among species within the *B. armata* clade in northwestern Mexico, and some evidence of niche non-equivalency between *B. berlandieri* and *B. dulcis*, the former of which is synonymized under *B. dulcis*.

**Conclusions:** Our findings have implications for the complex biogeographic history in Central America and Mexico, suggesting that sympatric speciation and dispersal are the predominant processes of species diversification. Future studies should include population-level sampling across the genus, along with morphological and ecological information, to assess distinctness among species and, particularly, levels of gene flow, in an integrative fashion.

## Introduction

The palms (family Arecaceae) are globally emblematic components of tropical and subtropical ecosystems (Uhl and Drandfield, 1987; Dransfield, Uhl, *et al.*, 2008; Asmussen *et al.*, 2006; Baker *et al.*, 2009; Baker and Dransfield, 2016; Balslev et al., 2016). Palms are notorious for having slow substitution rates among the monocot angiosperms, likely resulting from their large size and associated consequences for the inheritance of new mutations (Gaut *et al.*, 1992; Lanfear *et al.*, 2013; Barrett *et al.*, 2016a). Slow mutation rates equate to fewer informative molecular characters for phylogenetic analysis, and thus hamper our understanding of interspecific relationships in many clades. Resolved, strongly supported phylogenetic hypotheses form an essential basis for taxonomic research and subsequent inference of character evolution, biogeography, and niche evolution. Genome-scale datasets provide a solution, having become feasible to obtain with the widespread availability of high-throughput sequencing methods. Not surprisingly, palm systematists have begun to embrace phylogenomic approaches (Comer *et al.*, 2015; Barrett *et al.*, 2016a; Comer *et al.*, 2016; Barrett *et al.*, 2016b; Heyduk *et al.*, 2016; Bacon *et al.* in review).

The fan palm genus *Brahea* Endl. *ex* Mart. exemplifies the situation described above, and is among the most taxonomically problematic and poorly understood clades of American palms (subfamily Coryphoideae, tribe Trachycarpeae, within which the placement of *Brahea* is uncertain; Quero and Yáñez, 2000; Quero, 2000; Hodel, 2006; Dransfield et al., 2008; Bacon *et al.*, 2012; Barrett et al., 2015; Baker and Dransfield, 2016). The genus comprises 11 currently recognized species (Hodel, 2006; Govaerts et al., 2018), that grow in dry, often calcareous soils in the coastal and lower montane regions of Mexico and Central America extending to Nicaragua. *Brahea* sometimes occurs with *Sabal* and *Washingtonia*, two other fan palm genera. The costapalmate leaf blades (i.e. extension of petiole into the blade) distinguish *Sabal* from *Brahea. Washingtonia*, which occurs natively in Baja California and Sonora, Mexico and in isolated oases in California and Arizona (and perhaps Nevada?), USA, has a similar overall appearance to some *Brahea* species, but differs in its bright to dull green leaves, whereas co-occuring *Brahea* (*B. armata*, *B. brandegeei*) have blue-green or gray leaves. Florally *Brahea* has three carpels connate only in the styles while *Sabal* has three carpels connate throughout. *Brahea* has tubular inflorescence rachis bracts while those of *Washingtonia* split on one side, become pendulous and are sword shaped.

Endlicher (1837) first informally used the genus name *Brahea*, honoring the Danish astronomer Tycho Brahe, and Martius (1838) validly published it a year later. Watson (1880) subsequently described the genus *Erythea*, which contains some species currently recognized in *Brahea. Brahea* was originally distinguished as having unarmed petioles, relatively small fruits, and solitary flowers, whereas *Erythea* was distinguished by having armed petioles, larger fruits, and clustered flowers at the base of the inflorescence. Moore (1973) and Uhl and Dransfield (1987) recognized these genera as a single genus, *Brahea*, with two subgenera—*Brahea* and *Erythea.* More recently, Quero and Yáñez (2000), and subsequently Hodel (2006) have refined various names associated with *Brahea* into 12 and 11 species, respectively, based on morphological and ecological observations. While other treatments recognize nine species (e.g. Henderson et al. 1995), here we follow Hodel (2006, see also Hodel 2018) and Govaerts et al. (2018) the most recent and comprehensive treatments of the genus.

Subgenus *Erythea*, as currently circumscribed, contains seven species. *Brahea aculeata*, *B. armata*, and *B. brandegeei* are found in the arid, desert scrub regions of northwestern Mexico and Baja California, and are distinguished by having distinctly armed petioles. *Brahea edulis* is endemic to Guadalupe Island, located approximately 240 km west of Baja California, and is IUCN red-listed as endangered in the wild (EN, C1; Johnson, 1998). This species is distinguished by unarmed petioles and large fruit size (> 25 mm in diameter). *Brahea pimo* occurs in the western pine-oak sierras of Mexico, and *B. salvadorensis* is restricted to El Salvador and Guatemala. These two species have distinctive hairy scales on the petioles and adaxial leaf surfaces and hairy-tomentose flowers, differing mainly in the degree of the latter.

Subgenus *Brahea* contains the widespread and highly variable *B. dulcis* and *B. calcarea* from Mexico into Guatemala, Honduras, and Nicaragua. The subgenus is distinguished mainly by absence or reduction of petiolar teeth. *Brahea dulcis* has small teeth at the base of the petiole, and inflorescences usually shorter than the leaves, whereas *B. calcarea* has unarmed petioles and inflorescences exceeding the length of the leaves. *Brahea berlandieri*, which is currently synonymized with *B. dulcis* (Govaerts and Dransfield, 2005; Hodel 2006; Govaerts et al., 2018; Hodel, 2018), occurs in northeastern Mexico, and was originally distinguished by having shorter rachillae than *B. dulcis* by Quero and Yáñez (2000). Though synonymized, it is unclear whether *B. berlandieri* is distinct from *B. dulcis* or whether it is a local form of the latter in northeastern Mexico. The two dwarf species *B. decumbens* (montane, limestone soils in open, rocky sites) and *B. moorei* (limestone soils in montane oak forest understory) differ in that *B. decumbens* has blue-gray leaves and a branching, creeping trunk, whereas *B. moorei* has leaves green adaxially and chalky white abaxially, is solitary, and appears trunkless. Quero (2000) subsequently described *Brahea sarukhanii* from a narrow region of Nayarit and Jalisco in central-western Mexico, but this species is of uncertain affinity in that it displays characteristics of both subgenera, including small teeth near the base of the petiole and relatively smaller fruits. Hodel (2006) distinguishes this species from the widespread and co-occurring *B. dulcis* by the persistence of old leaf bases along the entire length of the trunk as opposed to only along the upper third of the trunk (*B. dulcis*).

Despite recent taxonomic progress, there has been no explicit phylogenetic analysis of relationships in *Brahea.* Furthermore, the existence of morphological intermediacy, environmental and local variation, and hybridization/introgression (e.g. Ramirez-Rodríguez *et al.*, 2011) have likely precluded a definitive understanding of taxonomy in this genus. *Brahea* presents striking variation in growth form, with some species being trees of tall-to-medium stature at one extreme, some of variable or intermediate height (e.g. *B pimo*, *B. aculeata*), and some having small, acaulescent, creeping, shrub-like, or cespitose growth forms (e.g. *B. decumbens*, *B. moorei*). The two dwarf species, *B. decumbens* and *B. moorei*, grow in sympatry in northeastern Mexico, calling into question whether these species are closely related, or if they represent convergent growth forms within the genus. Thus *Brahea* represents a highly appropriate system in which to test the hypothesis that height and DNA substitution rates are negatively correlated (sensu Lanfear et al., 2013).

Species of *Brahea* occupy a wide variety of environmental conditions across the geographic range of the genus, from northern Mexico to Nicaragua, where they often represent major ecosystem structural components, especially in desert and other xeric habitats (Uhl & Dransfield, 1987; *Henderson *et al.*, 1995*; Quero, 2000; Hodel, 2006; Dransfield, Uhl, *et al.*, 2008; Wehncke *et al.*, 2013). However, the biogeographic history of this clade is unknown, and likely complex, potentially having been shaped by vicariance, dispersal, and sympatric speciation across a geologically dynamic landscape (e.g. Sedlock *et al.*, 1993). The geological history of Mexico is also complex, with several major events having occurred over the last few tens of millions of years that could have potentially shaped distributions and species divergence in *Brahea.* These include the formation of: (1) the Mexican Transvolcanic Belt ca. 15 million years ago (mya), bisecting mainland Mexico to the north and south; (2) the opening of the Gulf of California, separating Baja California from western mainland Mexico ca. 7.2 mya; and (3) the formation of Guadalupe Island via volcanic activity also around 7.2 mya (Ferrari *et al.*, 2012; Bennett and Oskin, 2014; Dolby *et al.*, 2015). It is currently unknown if or how these processes influenced diversification and dispersal in *Brahea*.

In addition to the potential allopatric influence of geological processes in diversification, several species of *Brahea* have contemporary, overlapping ranges, and may exemplify cases of sympatric niche differentiation and speciation. For example, *B. armata* overlaps with *B. aculeata* and *B. brandegeei* in northwestern Mexico, while *B. dulcis* overlaps with *B. calcarea*, *B. pimo*, *B. moorei*, *B. sarukhanii*, *B. decumbens*, and the synonymous “*B. berlandieri*” on mainland Mexico. Species distribution models (SDMs) allow us to test for ecological scenarios potentially underlying species diversification (Nunes and Pearson, 2017). We can test for phylogenetic niche diversification vs. conservatism (Harvey and Pagel, 1991; reviewed in Pyron *et al.*, 2015) by assessing the equivalence of inferred niche space between species pairs relative to niche differences due to random chance. We expect to find closely-related species that occupy distinct niche spaces, often with overlapping ranges or in close proximity.

Here we use a combination of genome skimming (plastid and mitochondrial genomes, ribosomal DNA cistrons) and Sanger sequencing of single-copy nuclear introns to generate a phylogenomic dataset of approximately 400 kb. Our specific objectives are to: (1) provide resolution and support for species-level phylogenetic relationships within *Brahea* and to test the current subgeneric circumscription; (2) test the putative association between plant height and substitution rate in a phylogenetic context via ancestral state reconstruction; (3) infer divergence times and biogeographic history; and (4) test for niche differentiation among closely related, geographically overlapping species. This study has implications for Central American biogeography (more prominent roles for dispersal and sympatric speciation than for allopatric speciation via vicariance), patterns of ecologically driven species diversification (sympatric niche differentiation), growth form evolution, horticulture, and conservation of these ecologically important palms.

## Materials and Methods

### Phylogenetic analyses

#### Taxon sampling and DNA sequencing

We sampled all 11 species currently recognized in *Brahea* (Table 1, including *Washingtonia robusta* and *Chamaerops humilis* as outgroups; Hodel, 2006; Govaerts et al., 2018), extracted DNAs using the CTAB method (Doyle and Doyle, 1987), and used standard procedures for Illumina library preparation and sequencing [Supplementary Information, Methods]. We amplified nuclear intron regions, corresponding to the Malate Synthase (*MS*) gene; Serine/Threonine Protein Kinase genes *CISP4* and *CISP5*; and DNA-directed RNA Polymerase Subunit Beta gene (*RPB2*) [Supplementary Information, Methods].

**Table 1.** Voucher information and characteristics of high-throughput sequence datasets. HBG = Huntington Botanical Garden live collection; HNT = Huntington Botanical Garden Herbarium; CSLA = California State University, Los Angeles Herbarium; UCBG = UC, Berkeley Botanical Garden live collection; UC = UC, Berkeley & Jepson Herbaria; ‘Plastid,’ ‘rDNA,’ and ‘mt’ = mean coverage depth of the plastome, partial rDNA cistron (18S-ITS-26S), and mitochondrial genomes, respectively. ^a^Recognized as a synonym of *B. dulcis* (Hodel, 2006; Govaerts et al., 2018).

| Species | Voucher | Total # reads | Plastid | rDNA | mt | Sequencing |
| --- | --- | --- | --- | --- | --- | --- |
| *Chamaerops humilis* L. | HBG 2073; HNT 10471 | 8,677,266 | 96.6 | 508.8 | 10.7 | HiSeq2000 |
| *Washingtonia robusta*<br>H.Wendl. | CSLA Barrett 310 CSLA | 37,162,404 | 1,128.0 | 2,096.3 | 176 | HiSeq2000 |
| *Brahea aculeata*<br>(Brandege) H.E.Moore | HBG 16448; HNT 13042 | 5,529,618 | 175.2 | 2,716.6 | 18 | MiSeq |
| *Brahea armata* S.Watson | HBG 23437; HNT 13222 | 7,273,550 | 301.3 | 3,848.1 | 19.6 | MiSeq |
| *Brahea “berlandieri”*<br>Bartlett<sup>a</sup> | HBG 28812; HNT 1745 | 5,051,496 | 100.6 | 529.6 | 29.8 | MiSeq |
| *Brahea brandegeei* (Purpus)<br>H.E.Moore | HBG 89871; HNT 13043 | 9,873,046 | 281.2 | 4,185.6 | 16.5 | HiSeq2000 |
| *Brahea decumbens* Rzed. | HBG 35650; HNT 13038 | 4,013,662 | 36.2 | 276.2 | 9.5 | MiSeq |
| *Brahea dulcis* (Kunth) Mart. | HBG 23220; HNT 13044 | 15,361,806 | 638.2 | 1,772.7 | 66.1 | NextSeq500 |
| *Brahea edulis* H.Wendl. ex<br>S.Watson | UCBG 2003.0149;<br>1869963, UC 1971077 | 18,933,406 | 554.7 | 3,626.7 | 42.1 | HiSeq2500 |
| *Brahea moorei* L.H.Bailey<br>ex H.E.Moore | UCBG 88.0345; UC<br>197077 | 3,103,380 | 59.3 | 1,160.6 | 9.3 | MiSeq |
| *Brahea calcarea* Liebm.<br>(syn. *B. nitida* André) | HBG 27970; HNT 10472 | 12,865,552 | 494.2 | 871.2 | 30.1 | NextSeq500 |
| *Brahea pimo* Becc. | HBG 52194; HNT 13045 | 6,518,544 | 42.8 | 615.3 | 11.4 | MiSeq |
| *Brahea salvadorensis*<br>H.Wendl. ex Becc. | HBG 35754; HNT 1029 | 4,917,552 | 142.8 | 439.2 | 20.8 | MiSeq |
| *Brahea sarukhanii*<br>H.J.Quero | HBG 25151; HNT 10470 | 15,333,496 | 279.8 | 1,617.1 | 29 | NextSeq500 |

#### DNA assembly, alignment and phylogenetic analyses

We assembled plastid, mitochondrial, and rDNA contigs de novo in NOVOPlasty (version 2.6.3; Dierckxsens *et al.*, 2016), and extended contigs in GENEIOUS v.11 (Biomatters Ltd., New Zealand). We aligned plastomes in MAFFT v.1.33 (Katoh and Standley, 2013), and MUSCLE (Edgar, 2004), and removed positions containing gaps. We subjected both separate and concatenated alignments to phylogenetic analysis under Parsimony, Maximum Likelihood, Bayesian, and quartet-based methods. We conducted: Parsimony searches in TNT with 2,000 jackknife replicates (Goloboff *et al.*, 2008); Maximum Likelihood (ML) searches in RAxML v.8 under a GTRGAMMA model with 1,000 standard bootstrap replicates (Stamatakis, 2014); Bayesian analyses in MrBayes v.3.2.6 (Ronquist *et al.*, 2012) for 10^7^ generations (GTR+Gamma+I model) and with the first 25% as burn-in; and quartet-based methods in SVDquartets using both concatenated and coalescent models, each with 1,000 bootstrap replicates (Chifman and Kubatko, 2014; 2015) [details of these analyses can be found in Supplementary Information, Methods]. We quantified congruence and conflict among ML topologies based on plastid, mitochondrial, nuclear rDNA, and nuclear introns using Robinson-Foulds (RF) tree distances (Robinson and Foulds, 1979) in PAUP v.4.0 (Swofford, 2002).

### Growth form evolution

We reconstructed the evolution of growth form among species of *Brahea* by using maximum plant height following values in Henderson *et al.* (1995), Quero (2000), and Quero and Yáñez (2000). We used the R (R Core Development Team, 2014) packages ‘ape’ (Paradis, 2004) and ‘phytools’ (Revell, 2012) to infer ancestral values for height under a Maximum Likelihood Brownian Motion model, based on the combined, total evidence tree with branch lengths converted to ultrametric via non-parametric rate smoothing (Sanderson, 1997) in R. We computed root-to-tip GTR distances using the ‘vcv.phylo’ function on the non-smoothed phylogram from above, and regressed these on maximum height values using standard and phylogenetic regression via Independent Contrasts (Felsenstein, 1980; Garland *et al.*, 1992) in the R package ‘ape’. We tested for phylogenetic signal in both maximum height and GTR-based branch lengths from the combined RAxML tree using Pagel’s λ.

### Biogeographic analyses

#### Divergence time estimation

We estimated divergence times among species of *Brahea* in BEAST2 (Bouckaert *et al.*, 2014) under a Lognormal Relaxed Clock (Drummond *et al.*, 2006) and a GTR+Gamma+I substitution model with parameters estimated from the data. We added data from all *Brahea* species to the dataset of Couvreur *et al.* (2011), for a total of nine loci [see Supplementary Information, Methods for additional details]. We chose four calibration points with exponential priors, to be consistent with the analysis of Couvreur *et al.* (2011): *Sabalites* (Berry, 1914), to calibrate the stem node of Coryphoideae at [minimum age offset = 85.8 million years ago (mya), mean = 1.0]; *Mauritiidites* (Schrank, 1994) to calibrate Mauritiinae (minimum age offset = 65.0 mya, mean = 1.5); a *Cocos*-like fossil to calibrate subtribe Attaleinae (minimum age offset = 54.8 mya, mean = 2.0); and *Hyphaenae kapelmanii* (Pan et al., 2006) to calibrate the stem node of Hyphaeninae (minimum age offset = 27.0 mya, mean = 0.5). We also constrained the stem node of palms (i.e. palms + Dasypogonaceae) to have arisen between 110-120 mya (these dates were used as the 95% prior on a normal distribution with a mean of 115 mya). We ran BEAST2 for 2×10^8^ generations of the MCMC, sampling every 5×10^4^ generations, discarding the first 30% of samples as burn-in after a pre-burn-in period of 10^5^ samples. We verified stationarity via effective samples sizes >200 for each parameter in TRACER (Rambaut *et al.*, 2014), and also verified convergence of parameter values by combining results of three independent runs of BEAST2 from random starting seeds.

#### Ancestral range reconstruction

We used the chronogram from BEAST2 to infer ancestral ranges of *Brahea* species in BioGeoBEARS (Matzke, 2014). We chose six areas, corresponding to major physiographic regions of Mexico and Central America (e.g. Thayer, 1916; Riddle *et al.*, 2000; Bennett and Oskin, 2014): A) the Baja Californian Peninsula; B) northwestern mainland Mexico, containing the Sierra Madre Occidental, Sonoran Basin-and-Range, and Pacific Coastal Plain; C) northeastern Mexico, containing the Sierra Madre Occidental, the Great Plain, and the Northern Gulf Coast Plain; D) central/southern Mexico, containing the Mexican Transvolcanic Belt and the Southern Sierra Madre; E) southeastern Mexico and Central America; and F) Guadalupe Island. We used a time-stratified approach, with two time intervals (25.7-7.2 mya, and 7.199-0 mya), corresponding to the formation of Guadalupe Island and the Gulf of California (i.e. the separation of Baja California from mainland Mexico) approximately 7.2 million years ago (Bennett and Oskin, 2014). We used no constraints between directly adjacent areas (i.e. dispersal probabilities of 1.0), a dispersal probability of 0.5 for non-adjacent areas, and a probability of 0.1 for dispersal probability between Baja California and southeastern Mexico/Central America. For the first interval (25.7-7.2 mya), dispersal probability between Guadalupe Island and all other areas was set to zero, and then to 0.01 for the second interval (7.199-0 mya). Also in the second interval, we set dispersal probability at 0.1 between Baja California and northwestern mainland Mexico. We used the BayAreaLike+j model in BioGeoBEARS (Matzke *et al.*, 2014).

#### Species distribution modeling and ecological niche differentiation

We inferred species distribution models (SDM) to: (1) determine if the contemporary ranges of *Brahea* species can be modeled using available bioclimatic predictor variables; (2) test signal of niche divergence for selected pairs of *Brahea* species with overlapping ranges; (3) assess the degree to which SDMs can inform taxonomic boundaries in the group. We obtained occurrence data for all species included in this study via the Global Biodiversity Information Facility (GBIF.org, accessed 16 January 2018), followed by filtering both manually and using the ‘trim.occ’ R function of Nunes and Pearson (2017).

We produced SDMs using MaxEnt (version 3.4.1; Phillips *et al.*, 2017) through the R package Dismo (version 1.1-4; Hijmans *et al.*, 2017). Two environmental datasets were used for SDMs: the entire WorldClim dataset (version 2.0; Fick and Hijmans, 2017), and a Pearson correlation-filtered version, both at ten-minute resolution [see Supplementary Information, Methods for additional details]. We tested pairs of species in the *B. armata* clade, and between “*B. berlandieri*” and *B. dulcis* for niche equivalency (Warren *et al.*, 2008) and phylogenetic niche conservatism (Harvey and Pagel, 1991; see Pyron *et al.*, 2015). We used the R function ‘nicheEquivalency’ in the Dismo package to conduct Warren’s Identity test; and the ‘Random Translocation and Rotation’ (RTR) and the modified niche overlap (MO) metric of Nunes and Pearson (2017) to test for Phylogenetic Niche Conservatism (RTR and MO are hereafter collectively referred to as the RTR-MO). We adjusted our alpha level for comparisons made of species in the *B. armata* clade using Bonferroni correction (α_adjusted_ = 0.016).

## Results

### Datasets

Features of all molecular datasets included in this study are summarized in Table 2. The complete plastome alignment of 11 *Brahea* and two outgroup taxa (*Chamaerops*, *Washingtonia*) is 132,870 bp in length. Removal of all sites with gaps/missing data and ambiguity codes produced an alignment of 121,301 bp with 333 parsimony-informative sites. Four contiguous regions of the mitochondrial genome were included here (approximately 67, 91, 52, and 25 kb, respectively), totaling 261,650 bp when concatenated. After removing gaps and ambiguities as above, the total aligned length was 212,874 bp with 304 Parsimony-informative sites. Amplicons for *CISP5* showed clear evidence of double banding, and so this locus was excluded from further analysis. The total, combined dataset was 342,142 bp in length, with 756 PIC. Data are deposited under GenBank accession numbers XXXXXX-XXXXXX.

**Table 2.** Characteristics of the nuclear, mitochondrial, and plastid datasets for *Brahea.* ‘Total L’ = total length of each alignment; ‘post-filter L’ = length of each alignment after filtering; ‘# P-inf sites’ = the number of parsimony informative sites; ‘P-inf/1000bp’ = the number of parsimony informative sites per 1,000 bp of alignment; ‘#MPT, L’ = the number of most parsimonious trees and their length in steps (i.e. number of nucleotide changes); ‘model’ = the best fit model for each alignment based on the corrected Akaike Information Criterion.’

| Locus | total L | post-filter L | # P-inf sites | P-inf/1000bp | #MPT; L | 837<br>model |
| --- | --- | --- | --- | --- | --- | --- |
| *MS* | 601 | n/a | 18 | 30.0 | 9, 41 | T92+G |
| *CISP4* | 664 | n/a | 22 | 33.1 | 3, 60 | GTR+G |
| *RPB2* | 701 | n/a | 19 | 27.1 | 1, 47 | HKY+G |
| **rDNA** | 6,180 | n/a | 60 | 9.7 | 3, 218 | TN93+G |
| **Total nrDNA** | 8,146 | 7,967 | 119 | 14.6 | 1, 384 | TN93+G |
| **mtDNA** | 261,650 | 212,874 | 304 | 1.2 | 1, 1,164 | GTR+G+I |
| **Plastome** | 132,870 | 121,301 | 333 | 2.5 | 1, 938 | HKY+G+I |
| **Combined** | 402,666 | 342,142 | 756 | 2.2 | 1, 2,490 | GTR+G+I |

### Phylogenetic analyses

#### Plastomes

Parsimony, Maximum Likelihood, and Bayesian analyses of whole aligned plastomes, with one copy of the IR removed, yield highly resolved and strongly supported relationships [Fig. 1A; Supplementary Information, Fig. S1]. *Brahea* is supported as monophyletic (Parsimony Jackknife = 100, ML Bootstrap = 100, Bayesian Posterior Probability = 1.0; denoted as ‘100, 100, 1.0’ hereafter), within which there are two strongly supported principal clades. The first of these (100, 100, 1.0) contains *B. decumbens*, *B. calcarea*, *B. dulcis*, and *B. berlanderi* (hereafter referred to as the ‘*B. dulcis* clade’). The latter two accessions share a close relationship (89, 94, 1.0), but relationships among *B. decumbens*, *B. calcarea*, and the clade of (*B. berlanderi*, *B. dulcis*) are unresolved. The next clade contains all remaining *Brahea* (100, 98, 1.0), and is further divided into two clades within which all relationships are strongly supported (≥97, 1.0). The dwarf species *Brahea moorei* is sister to *B. sarukhanii* (100, 100, 1.0), while *B. pimo* is sister to *B. salvadorensis* (100, 100, 1.0), and these two clades are sister to one another (100, 100, 1.0; hereafter referred to as the *B. pimo* clade). Sister to this is a clade composed of (*B. armata*, *B. aculeata*; 90, 93, 1.0), successively sister to *B. brandegeei* (98, 97, 1.0), and *B. edulis* (100, 100, 1.0), hereafter referred to as the ‘*B. armata* clade.’

**Fig. 1.**
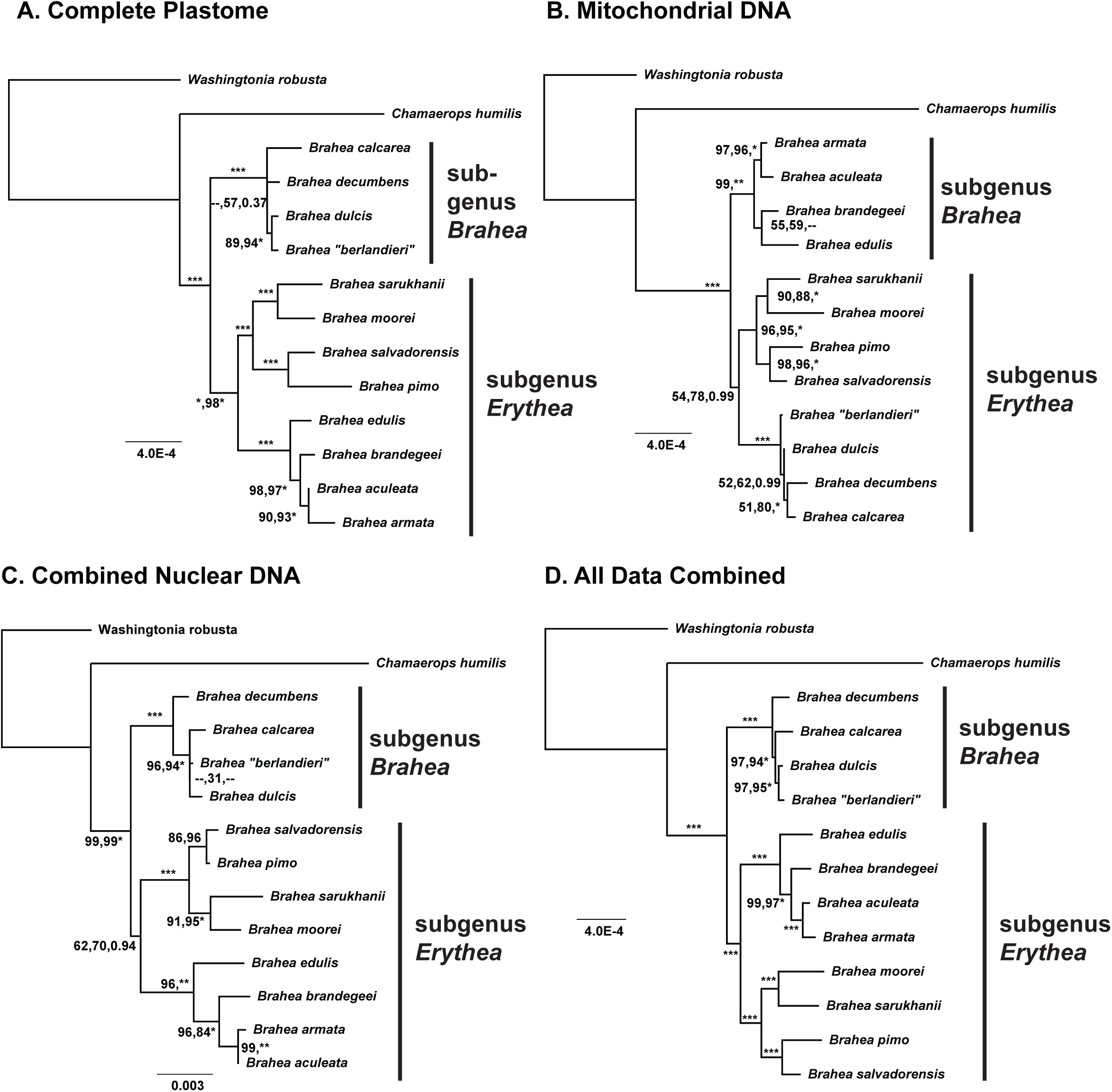
Maximum Likelihood trees based on plastid (**A**), mitochondrial (**B**), nuclear (**C**), and combined (**D**) data. Numbers adjacent to branches are Parsimony Jackknife, Maximum Likelihood Bootstrap, and Bayesian posterior probabilities. ‘^∗^’ = 100% support (or posterior probability = 1 for Bayesian Analysis). ‘‐‐’ = differing topology from ML tree for Parsimony or Bayesian Analysis. Scale bar = substitutions/site.

#### Mitochondrial DNA

Relationships overall are less strongly supported for mtDNA than for plastomes (Fig. 1B). The three primary clades recovered from the plastome data are also recovered for mtDNA. A notable difference is the placement of the *B. dulcis* clade as sister to the *B. pimo* clade, collectively sister to the *B. armata* clade, but this relationship is weakly supported (54, 78, 0.99). Many sister relationships are identical to those from the plastome and are well supported (e.g. *aculeata+armata; moorei*+*sarukhanii; pimo*+*salvadorensis*), while remaining sister relationships among species that differ from those of the plastome receive weak support.

#### Nuclear DNA

Relationships based on nuclear ribosomal DNA (rDNA) are highly identical to those based on plastomes, albeit with lower support overall, and differing only in the placement of *B. decumbens* [Supplementary Information, Fig. S1]. Both Malate Synthase (*MS*) and *CISP4* recover the same ‘deep’ tree structure among the three principal clades, but with weak support individually [Supplementary Information, Fig. S1]. *RPB2* displays several relationships not observed for any other locus, but most have weak support; e.g., *B. moorei* and *B. brandegeei* group in a clade of *B. dulcis*, *B. berlandieri*, and *B. calcarea* [Supplementary Information, Fig. S1].

#### Combined analyses

Combined nuclear data (rDNA, *MS*, *CISP4*, and *RPB2*) yield a highly supported topology similar to that based on plastomes, differing only in the placement of *B. decumbens* (Fig. 1C). Analysis of the concatenated dataset from all three genomes recovers a highly supported tree (Fig. 1D; all support values >94). These relationships differ from those of the plastome only in the placement of *B. decumbens* as sister to (*B. calcarea*, (*B. dulcis*, *B. berlandieri*)). Analysis in SVDquartets (specifying one tree for all sites) yields an identical topology to those based on Parsimony, Maximum Likelihood, and Bayesian Analysis [Supplementary Information, Fig. S2]. In this SVDquartets analysis, a topology of (*B. edulis*, (*B. brandegeei*, (*B. armata*, *B. aculeata*))) is recovered with 100% Bootstrap support for all relationships; by contrast, a topology of (*B. brandegeei*, (*B. edulis*, (*B. armata*, *B. aculeata*))) is recovered under the multispecies coalescent model, with only 39% Bootstrap support for the placement of *B. edulis* as sister to (*B. armata*, *B. aculeata*). A second coalescent-model analysis in SVDquartets, specifying the ‘Erik+2’ parameter, places *B. edulis* as sister to *B. brandegeei* with low support (Bootstrap = 54), and these are sister to (*B. armata*, *B. aculeata*). Thus, there is little support in coalescent analyses for the relative placement of *B. edulis* and *B. brandegeei* within the *B. armata* clade.

Robinson-Foulds tree distances on average were greatest between the *CISP4* topology and all other topologies [Supplementary Information, Table S1]. Out of the three largest single-genome alignments (i.e. combined nrDNA, mtDNA, and plastomes), the combined nuclear tree had the lowest RF distance to the total combined tree (2×RF = 0), followed by the plastome tree (2×RF = 2), and the mtDNA tree (2×RF = 6).

### Growth form evolution

Maximum height in *Brahea* ranges from 0.4 m in *B. moorei* to 15 m in *B. armata.* The tallest species occupy the *B. armata* clade [ancestral value = 10.63 ± 2.08 m (i.e. ± one standard deviation)], while the shortest species occupy the *B. pimo* clade (ancestral value = 5.32 ± 2.31 m) (Fig. 4). Both of these clades comprise subgenus *Erythea* sensu Quero and Yáñez (2000) (ancestral value = 6.95 ± 2.63 m). The ancestor of subgenus *Brahea* is estimated to have had a maximum height of 6.37 ± 1.85 m. Maximum height shows a strong relationship with root-to-tip branch lengths, which range from 0.000469 substitutions·site^-1^ (s·s^-1^) in *B. dulcis* to 0.000803 s·s^1^) in *B. moorei* (F = 32.94, df = 11, *p* = 0.00013). However, this relationship is non-significant when correcting for phylogenetic relationships via Independent Contrasts (F_pic_ = 0.002, df = 10, *p* = 0.96), suggesting that any relationship between these traits is better explained by shared ancestry. Pagel’s λ, a measure of phylogenetic signal, was significantly different than a model of λ = 0 for root-to-tip branch lengths (λ = 1.06, P = 0.0009), but not for maximum height (λ = 0.61, P = 0.15). The two shortest species likely evolved from taller ancestors: maximum height in *B. decumbens* is 2.5 m vs. the ancestral value of 6.37 ± 1.85 m for the *B. dulcis* clade, while in *B. moorei* maximum height is 0.4 m vs. the ancestral values of 5.14 ± 2.17 m for (*B. moorei*, *B. sarukhanii*).

### Biogeographic analyses

#### Divergence time estimation

The estimated stem-node age of *Brahea* is 25.6 million years ago (mya), with a 95% highest posterior density (HPD) of 14.8-35.5 mya, while the crown age of *Brahea* is estimated to be 15.6 mya (9.1-22.2) (Fig. 2). Estimates of the crown radiations of the three principal clades of *Brahea* are: 5.5 mya (2.1-9.8) for the *B. armata* clade; 7.6 mya (3.3-12.0) for the *B. pimo* clade; and 6.2 mya (2.2-11.1) for the *B. dulcis* clade. *Brahea armata* and *B. aculeata* likely diverged relatively recently (1.36 mya, 0.05-2.6), as did *B. dulcis* and *B. berlandieri* (0.5, 0.00-1.4).

**Fig. 2.**
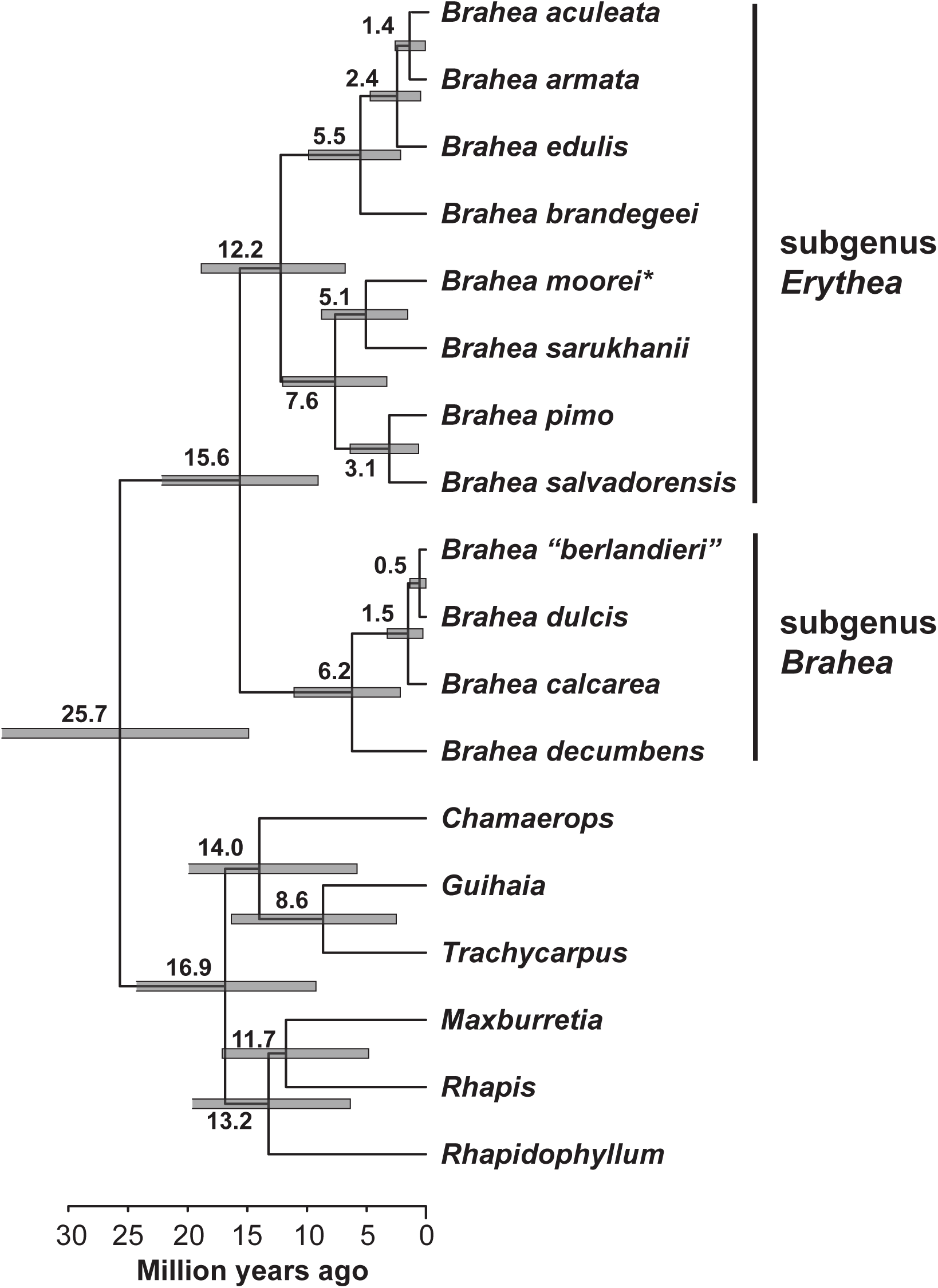
Divergence time estimates for *Brahea* based on four fossil calibrations. Scale bar = millions of years before present. Gray bars on nodes = 95% highest posterior density of divergence time estimates; numbers at nodes are mean divergence time estimates. “^∗^” indicates that *B. moorei* is placed in subgenus *Erythea* based on our phylogenetic results, and not in subgenus *Brahea* as in previous treatments.

#### Ancestral range reconstruction

We inferred an ancestral range of AB for the *B. armata* clade (Fig. 3; Baja California + northwestern mainland Mexico), and all nodes within this clade are estimated to share the same ancestral range, all with high likelihood percentages for the BayAreaLike+j model (L_%_ > 0.95). However, L_%_ is substantially lower when the ‘j’ parameter is excluded. *Brahea edulis* shows a high L_%_ of having arisen via founder event speciation on Guadalupe Island when including the ‘j’ parameter (jump dispersal). Ancestral ranges are more ambiguous in the *B. pimo* clade (Fig. 3), with estimates of C (northeastern Mexico) or D (central/southern Mexico) for the ancestral ranges of *B. moorei* and *B. sarukhanii*, and D or E (southeastern Mexico/Central America) as the ancestral range of *B. pimo* and *B. salvadorensis.* An ancestral range of BDE or BCDE (northwestern mainland Mexico, central/southern Mexico, southeastern Mexico/Central America) is inferred for the common ancestor of the *B. dulcis* clade (L_%_ = 0.60 including ‘j’; L_%_ = 0.10 excluding ‘j’). The ancestral range of *Brahea* as a genus is equivocal based on the current analysis (BDE L_%_ = 0.18 including the ‘j’ parameter; ABCD L_%_ = 0.04 excluding ‘j’).

**Fig. 3.**
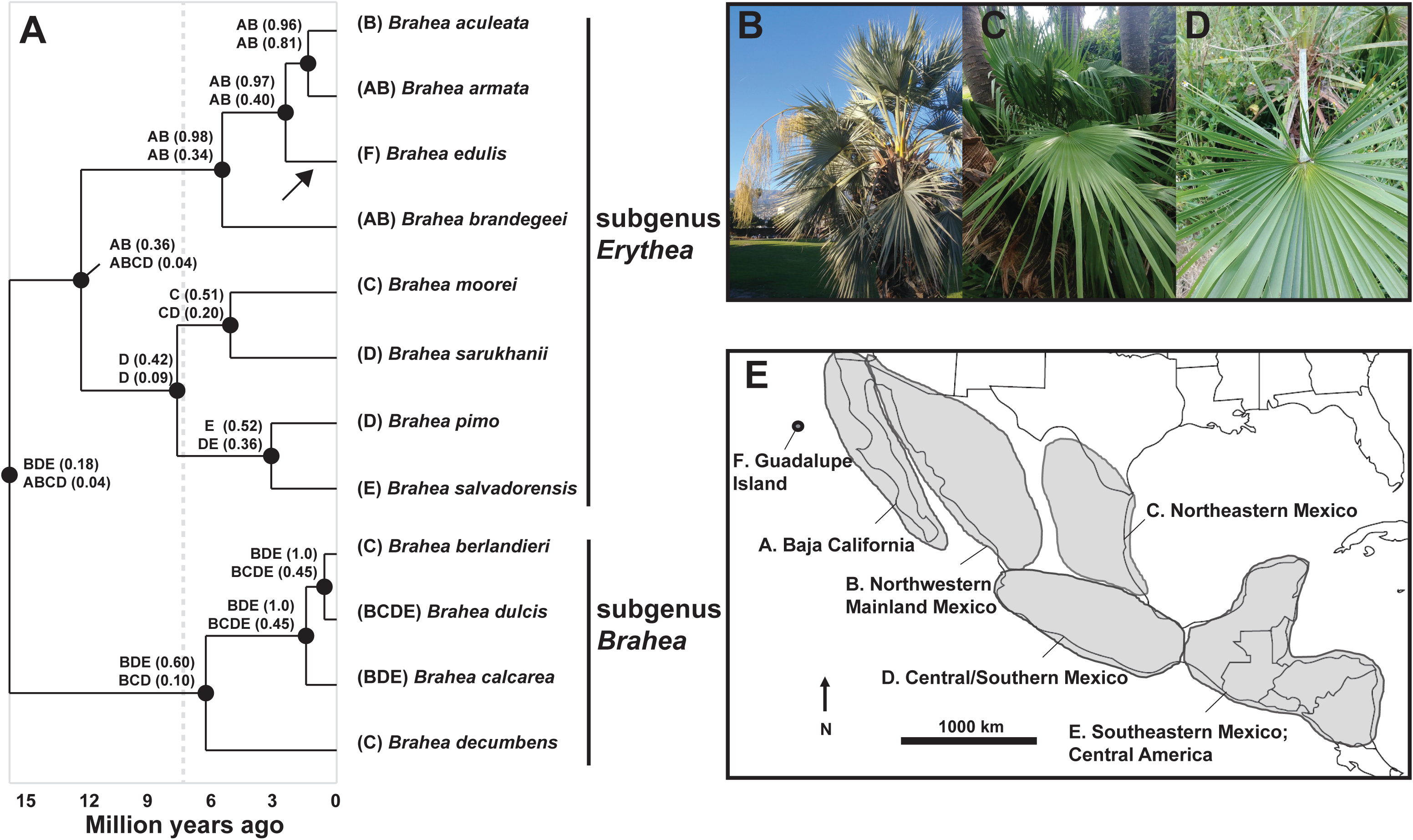
Biogeographic reconstruction of ancestral ranges for *Brahea* under BayAreaLike and BayAreaLike+j models in BioGeoBEARS. **A**. Chronogram with most likely ancestral ranges. Numbers adjacent to pie charts indicate the likelihood of the most likely ancestral range (top BayAreaLike, bottom, BayAreaLike+j). Dotted line indicates the approximate dates of origin of Guadalupe Island and the formation of the Gulf of California (7.2 mya). **B**. *Brahea armata* (Santa Barbara, California, USA). **C**. *Brahea dulcis* (Fairchild Tropical Botanical Gardens, Coral Gables, Florida, USA). **D**. *Brahea sarukhanii* (Montgomery Botanical Center, Coral Gables, Florida, USA). Photos: C. Barrett. **E**. Map of Mexico and Central America displaying the six different areas chosen to represent of the range of *Brahea*: A. Baja California; B. Northwestern Mainland Mexico; C. Northeastern Mexico; D. Central/Southern Mexico; E. Southeastern Mexico/Central America; F. Guadalupe Island. Geological events (circles): “**I**.” Formation of the Mexican Transvolcanic Belt (15 mya onward); “**II**.” Opening of the Gulf of California (7.2 mya onward); “**III**.” Formation of Guadalupe Island (7.2 mya).

#### Species distribution modeling and ecological niche differentiation

We rejected equivalency of SDMs for all comparisons between species pairs of interest (Table 3). RTR-MO tests identified signal of niche divergence between *B. brandegeei* and *B. aculeata* (P = 0.00983, Supplementary Information, Fig. S3], as well as between *B. aculeata* and *B. armata* [P = 0.00031, Supplementary Information, Fig. S3). Niche space overlap between *B. brandegeei* and *armata* falls below the established 5% CI of the null distribution, but is not significantly different from the null post-Bonferroni correction (P = 0.0374; α_adjusted_ crusted critical value = 0.016). The RTR-MO test also failed to reject that the observed overlap between *B. berlandieri* and *dulcis* significantly differed from the null distribution (P = 0.216).

**Table 3.** Results of Warren’s Identity Test (*I*) run for 1,000 replicates using both the WorldClim and Pearson-Filtered data sets (α ≤ 0.0166 for the *B. armata* clade, post-Bonferroni correction).

| Species Comparison | *I* statistic | P-Value |
| --- | --- | --- |
| *B. armata* vs. *aculeata* | 0.249853356 | $1.44 \times 10^{-45}$ |
| *B. armata* vs. *brandegeei* | 0.264126907 | $5.00 \times 10^{-23}$ |
| *B. aculeata* vs. *brandegeei* | 0.126009409 | $6.02 \times 10^{-15}$ |
| *B. dulcis* vs. “*berlandieri*” | 0.179367929 | $7.09 \times 10^{-64}$ |

We ultimately excluded approximately half of the GBIF localities, after manually and ecologically filtering outliers resulting from: taxonomic uncertainty, obvious cultivation out of the native range, or a lack of georeferencing/coordinate certainty [Supplementary Information, Table S2]. We identified no major differences between published species ranges and the extents of our filtered localities or replicated SDMs, with the exceptions of *B. calcarea* and *B. decumbens*, based on visual cross-validation using range maps of Henderson *et al.* (1995).

Analyses using our Pearson-filtered predictor dataset conservatively resulted in more overestimation of SDM extent, relative to those using the entire Worldclim dataset. The SDMs and associated downstream tests we report were based on the Pearson-filtered dataset [Supplementary Information, Table S2], in order to minimize false positives (which can be attributed to uncertainties associated with the use of publicly available data), and model overfitting.

MaxEnt analyses accurately predict the present-day extent of *Brahea* species using Bioclim variables [Supplementary Information, Table S3]. Models did not appreciably vary between MaxEnt replicates, and the predictor variables that contributed the most to each SDM were found to be both suitable and stable, as judged by marginal and SDM-specific response curves [see Supplementary Information, Link 1]. Area under the curve of the receiver operator characteristic, and the percent contributions of predictor variables to each SDM is reported in Supplementary Information, Table S4. The majority of variation observed between SDM extents was attributable to over-prediction of areas of low occurrence probability (< 0.4). Unique combinations of environmental predictor variables influenced the SDM of each species [Supplementary Information, Table S4], but some clade-specific predictor variable influences are evident. Over-prediction of occurrence and coarser resolution of predictor variables here result in more conservative results (though this may seem counterintuitive), since each measures observed overlap of estimated niche space. The extent of SDMs for *B. nitida* and *B. decumbens* were strongly over-predicted relative to published ranges, including many points outside of the predicted range, and a small area of occurrence, respectively. We thus chose not to include these two species in tests of niche equivalency or phylogenetic niche conservatism in the *B. dulcis* clade, considering the perceived low quality of SDMs inferred for *B. nitida* and *B. decumbens*.

## Discussion

Palm systematics has been a challenge due to a combination of morphological homoplasy and extremely slow plastid DNA substitution rates (Uhl and Dransfield, 1987; Uhl *et al.*, 1995; Gaut *et al.*, 1992; Barrett *et al.*, 2016a). Because of this, palm systematists have turned to genome-scale datasets, as these greatly increase the number of informative characters available for phylogenetic analyses (Barrett *et al.*, 2016a; Comer *et al.*, 2015; 2016; Barrett *et al.*, 2016b; Heyduk et al, 2016). Here we have sequenced over 400 kb of DNA from all three plant genomes, largely resolving and providing support for relationships among currently known species of *Brahea.* Furthermore, this study is the first to estimate divergence times, ancestral ranges, ancestral growth forms, and ecological niche space for this highly variable, taxonomically complex, and ecologically/economically important (e.g. Pulido and Coronel-Ortega, 2015) group of American palms.

### Phylogenetic Analyses

Despite the large amount of data generated per species of *Brahea*, we recovered fewer than 1,000 total informative positions from three genomes for our combined analysis. Yet, this number was sufficient to provide highly supported relationships among species of *Brahea.* Information content varies from an average of 1.2 informative sites/1,000 bp in mtDNA to 33.1 sites/1,000 bp in nuclear *CISP4* (Table 2), underscoring the importance and power of including nuclear DNA in phylogenetic analyses of palms (e.g. Heyduk *et al.*, 2016).

Concatenated analyses yield high support for a single set of relationships, but coalescent based analyses in SVDquartets differ in the placement of *B. edulis/B. brandegeei* of the *B. armata* clade, with little to no support [Fig. 1; Supplementary information, Fig. S1 and S2]. Differing relationships based on coalescent methods indicate potential conflict among datasets, suggesting that incomplete lineage sorting, gene flow, or both may influence the recovered relationships in *Brahea.* This finding underscores the need to include many nuclear loci and multiple representatives of each putative species, such that scenarios of ILS and gene flow can be modeled and explicitly differentiated. Ways to accomplish this include examining the distribution of gene trees via concordance factors (e.g. Ané, 2007; Crowl *et al.*, 2017), or using methods that specifically incorporate gene flow (e.g. Jackson *et al.*, 2017; Morales *et al.*, 2017).

The two principal clades recovered here correspond closely to the subgenera *Brahea* and *Erythea* recognized by Moore (1973), Uhl and Dransfield (1987), Quero and Yáñez (2000), and Hodel (2006). *Brahea moorei*, a dwarf species from northeastern Mexico, is strongly supported as being a member of subgenus *Erythea*, in contrast to its placement in subgenus *Brahea* by Quero and Yáñez (2000). The more recently described *Brahea sarukhanii* (Quero, 2000), hitherto unplaced among other species of *Brahea*, shows a highly supported sister relationship to *B. moorei*, despite its caulescent habit and restricted distribution in Jalisco/Nayarit of western central Mexico.

Only two other studies to date have included multiple representatives of *Brahea.* Bacon *et al.* (2012) addressed biogeographic questions in the diverse fan palm tribe Trachycarpeae, which includes *Brahea.* Their analysis included four species of *Brahea*; three accessions of *B. dulcis* were sister to a clade of (*B. aculeata*, (*B. armata*, *B. brandegeei*)). Though sampling is limited in that study for *Brahea*, there is some agreement with the current study in that *B. dulcis* is separated from the *B. armata* clade. In that study, *B. brandegeei* is sister to *B. armata*, as opposed to *B. aculeata* being sister to *B. armata* in the current study, though this relationship is only moderately supported in the former. Klimova *et al.* (2017) used population-level sampling to address biogeographic questions in *Washingtonia robusta*, *W. filifera*, *Brahea armata*, *B. brandegeei*, and *B. edulis* in Baja California and Guadalupe Island; they also included a single specimen identified by the authors as ‘*B. elegans*’ (synonymized with *B. armata*) from eastern Sonora. Sequencing over 2 kb of plastid DNA yielded no variation among *B. armata*, *B. brandegeei*, and *B. edulis*, but their sample identified as *B. elegans* differed by five plastid substitutions, suggesting this sample might be more closely allied with another species of *Brahea* (Klimova *et al.*, 2017). Nuclear DNA sequencing yielded evidence of haplotype sharing among *B. armata*, *B. brandegeei*, and *B. elegans*, but a distinct haplotype for *B. edulis* from Guadalupe Island. Their findings further suggest a complex evolutionary history of *Brahea* in the region, and highlight the possibility of gene flow within the *B. armata* complex, though this cannot be distinguished from incomplete lineage sorting based on their data. Future sampling efforts should include numerous high-variation nuclear markers, complete or nearly complete organellar genomes, and a comprehensive sampling of multiple individuals from all currently recognized species of *Brahea* across their respective geographic ranges.

### Growth form evolution

Maximum Likelihood reconstruction of plant height suggests that the ancestor of *Brahea* was a medium-statured tree (6.85 ± 3.04 m). Estimated ancestral heights change little moving from the root ancestor in Fig. 4 to the ancestors of the *B. pimo* and *B. dulcis* clades, but increase dramatically in the ancestor of the *B. armata* clade, which contains all of the tallest species of *Brahea.* Interestingly, the two shortest species, *B. moorei* and *B. decumbens*, each likely evolved from medium-statured, caulescent ancestors, from which they diverged ca. 5.1 and 6.2 mya, respectively. The latter findings are likely responsible for the non-significance of phylogenetic signal in maximum plant height based on Pagel’s λ. Dwarf or small-statured palm species are often closely related to medium or tall-statured species (e.g. the dwarf species *Sabal minor* and *S. etonia*; *Zona, 1990*; Henderson *et al.*, 1995; Heyduk *et al.*, 2016), suggesting evolution in height differences can be rapid, as is the case between *B. moorei* and *B. sarukhanii*, and *B. decumbens* and taller members of the *B. dulcis* clade. *Brahea moorei* and *B. decumbens* are often referred to as dwarf species, yet they differ in some key ways. First, *B. moorei* has a short, solitary, rhizomatous, subterranean trunk (more rarely, the trunk can be aboveground), giving the plant a low-growing, shrub-like appearance (Henderson *et al.*, 1995; Hodel, 2006). *Brahea decumbens*, on the other hand, has a short, clustered, above-ground trunk, and often assumes a creeping habit (hence the species name ‘*decumbens*’), but is sometimes erect. In fact, as Hodel (2006) observed, there are many intermediate forms between *B. decumbens* and the widespread *B. dulcis*, suggesting environmental or local variation, and quite possibly gene flow among these species.

**Fig. 4.**
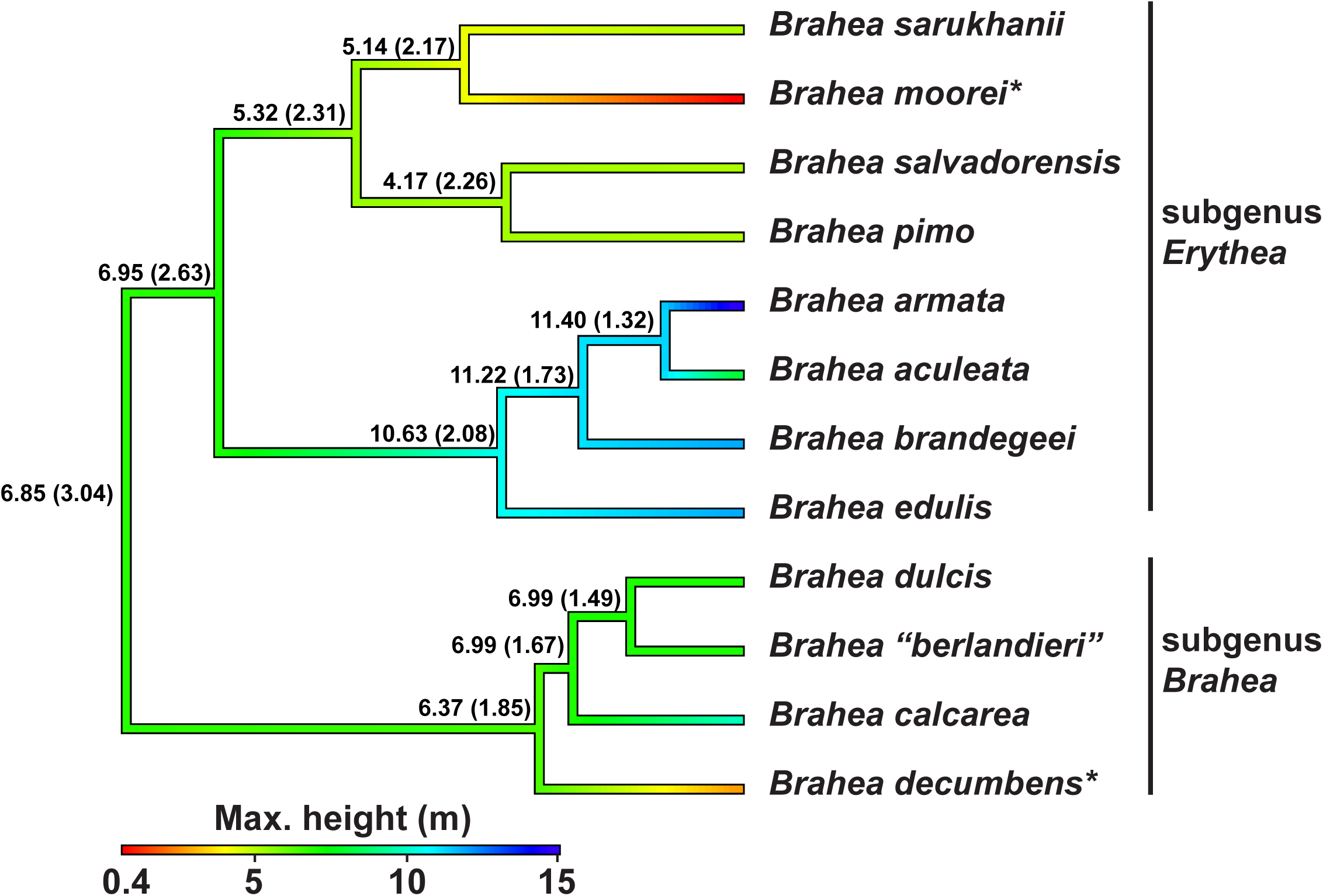
ML ancestral state reconstruction of maximum height among species of *Brahea*, based on a Brownian Motion model in the R packages ‘APE’ and ‘PhyTools.” Numbers at tips are maximum height values (in meters), and at nodes these are ancestral estimates of maximum height, with one standard error in parentheses. “^∗^” indicates the two dwarf forms.

A recent study across flowering plants demonstrated a large-scale relationship between plant height and substitution rates, when controlling for other factors such as species richness and latitude (Lanfear *et al.*, 2013), in which taller plants tend to have slower rates of substitution. This situation is pronounced in palms, which display some of the lowest substitution rates among monocots, and are unequivocally the tallest of the monocots (Barrett *et al.*, 2016a). Here, although these patterns exist at broad taxonomic scales (e.g. across angiosperm orders and families), it is unknown whether they also exist at finer taxonomic scales. Thus, we used *Brahea* as a case to address whether this pattern holds at finer taxonomic levels. Though there is an apparent negative correlation between substitution rate and height, this relationship is better explained by common ancestry, as indicated by Phylogenetically Independent Contrasts and significant phylogenetic signal in branch lengths based on Pagel’s λ. The relationship between substitution rates and height in plants has been explained by the ‘rate of mitosis’ hypothesis (Lanfear *et al.*, 2013), in which taller plants typically experience a slow-down of mitosis as they reach maximum height, and thus fewer potential mutations are passed on via reproductive tissues relative to the situation in shorter species. However, both *B. moorei* and *B. decumbens*, the two shortest *Brahea* species, are also extremely slow-growing (Hodel, 2006; S. Lahmeyer, personal observation), which may obscure any potential effect of plant height on heritable substitution rates in the genus, if such a relationship exists. Alternatively, the relationship may not exist at such low taxonomic levels, in which tall vs. short forms have recently evolved, and differences in substitution rates may potentially be determined by a number of factors differing on a taxon-by-taxon basis, as suggested by Barrett *et al.* (2016a).

### Biogeographic analyses

#### Divergence time estimates and ancestral range reconstruction

The stem and crown age estimates of *Brahea* (approximately 25.7 and 15.6 mya, respectively; Fig. 2) correspond closely with those in a previous study that included some accessions of *Brahea* (Bacon *et al.*, 2012). Our estimates suggest that ancestral forms of *Brahea* were present during some of the major geological events in Mexico, including the formation of the Transvolcanic Belt in Central Mexico (starting in the mid-Miocene, ca. 15 mya; Ferrari *et al.*, 2012), the Gulf of California (ca. 7.2 mya; Bennett and Oskin, 2014), and Guadalupe Island (also 7.2 mya), all around the same time (Dolby *et al.*, 2015).

The earliest divergence in *Brahea* corresponds approximately with the onset of formation of the Transvolcanic Belt in Mexico ca. 15 mya, though the estimated ancestral range of the common ancestor of all *Brahea* is equivocal based on our analysis in BioGeoBears (Fig. 3). Thus it is unclear if this geological process may have influenced diversification in *Brahea.* The most likely ancestral range for subgenus *Erythea* is Baja California and northwestern Mexico, but with low confidence. Even so, it is clear that geography has played a role in diversification in *Brahea*, with the *B. armata* clade concentrated in the northwest, members of the *B. pimo* clade occupying central, northeastern, and southeastern Mexico/Central America, and the *B. dulcis* clade occupying all areas but Baja California and Guadalupe Island.

The radiation of the *B. armata* clade began an estimated 5.51 mya (HPD = 2.13-9.84), which overlaps with the formation of Guadalupe Island and the Gulf of California (Bennett and Oskin, 2014). Jump dispersal (i.e. founder event speciation) can be attributed to the origin of *B. edulis* on Guadalupe Island. Based on Fig. 3, it is likely that the ancestor of *B. edulis* colonized Guadalupe Island soon after its formation (ca. 7.2 mya), and has since existed in isolation. The exact position of *B. edulis* varies among analyses here, either as sister to the remaining members of the *B. armata* clade, or as sister to (*B. armata*, *B. aculeata*) (Figs. 2, 3). Thus, it is unknown whether dispersal of the ancestral form of the *B. armata* clade occurred before or after the evolution of *B. brandegeei* in Baja California.

The ancestral range for the *B. armata* clade (Baja California and Northwestern Mexico) suggests the following sequence of biogeographic events were possible based on interpretation of Fig. 3: (1) sympatric range-copying in the ancestor of *B. brandegeei* and the remaining members of the *B. armata* clade; (2) founder event speciation as a result of jump dispersal of the ancestor of *B. edulis* to Guadalupe Island; and (3) subset/sympatric speciation of *B. aculeata* (northwestern Mexico) and *B. armata* (Baja California and northwestern Mexico). These findings contrast with those in other studies suggesting that the formation of the Gulf of California represents a major driver of allopatric speciation via vicariance (e.g. Riddle *et al.*, 2000). Instead, most speciation events in this clade likely occurred after the separation of Baja California from western mainland Mexico, suggesting roles for sympatric speciation or dispersal, especially in the case of *B. edulis*.

It is more difficult to interpret the estimated biogeographic history of the *B. pimo* clade, due to the equivocal likelihood percentage of its ancestral range in both models (Fig. 3). However, all speciation events in this clade potentially overlap with the ongoing formation of the Transvolcanic Belt in central Mexico, and thus it is unknown whether the later stages of volcanic uplift would have contributed to, for example, the divergence of *B. moorei* in northeastern Mexico and *B. sarukhanii* in central/western Mexico. Our results suggest a larger role for dispersal to neighboring areas than for vicariance and allopatric speciation due the formation of the Transvolcanic Belt (Fig. 3).

DEC-like models, and especially those carrying the ‘j’ parameter have been criticized recently, in that they fail to properly model cladogenetic events by preferentially biasing analyses towards cladogenesis (as opposed to anagenetic processes), and artificially inflating conclusions of jump dispersal/founder event speciation (Ree and Sanmartín, 2018). Therefore, we interpret our findings cautiously, and conclude that our proposed ancestral ranges for the species of *Brahea* are largely equivocal. The biogeographic history of Mexico and Central America is complex, and our ability to reconstruct the history of *Brahea* is limited in this case. Additional sampling within species, improved taxonomic delimitations, and more appropriate models including anagenetic and cladogenetic change in a time-dependent fashion while incorporating phylogenetic uncertainty (e.g. ClaSSE-type models; Fitzjohn, 2012) may help improve estimates of ancestral ranges in *Brahea*.

#### Species distribution models and ecological niche differentiation

Here we tested for signal of ecological diversification among selected species pairs within the *B. armata* and *B. dulcis* clades. These clades contain the youngest nodes in our chronogram, display overlapping species ranges in some cases, and contain taxonomic boundaries that remain somewhat uncertain (e.g. in the *B. dulcis* clade). The recent origin of *Brahea aculeata* and *armata* may be the result of sympatric ecological diversification in novel habitats. The respective SDMs of *Brahea aculeata* and *armata* are unique, even though their geographic ranges overlap. *Brahea brandegeei* and *armata* are distributed in Baja California and northwestern mainland Mexico, while *B. aculeata* is found only in northwestern mainland Mexico (Henderson *et al.*, 1995; Hodel, 2006). However, filtered GBIF records did not include *B. brandegeei* localities from northwestern mainland Mexico, an area listed by Henderson *et al.* (1995) to be part of this species’ range. Our inferences of SDMs and phylogenetic relationships between these three species suggest that the northern range and possibly the taller height of *B. armata*, and an eastern range extent and preference for more upland habitats by *B. aculeata*, may be evidence of relatively recent niche divergence. These results provide an additional line of evidence supporting our findings of sympatric speciation in the diversification of the *B. armata* clade in northwestern Mexico.

Recent taxonomic treatments place *B.* “*berlandieri*” as a synonym of the widespread and variable *B. dulcis* (e.g. Henderson et al., 1995; Hodel, 2006). Our inferred SDMs provide some evidence for the recognition of northeastern populations of *B. dulcis* corresponding to *B. berlandieri* (currently known as the former) as a distinct entity, raising the possibility that *B. berlandieri* could in fact be a separate species. Despite their estimated recent divergence time, their respective SDMs are significantly non-equivalent (0.18, P = 7.09 × 10^-64^). However, the failure to distinguish this difference from a null distribution argues against niche divergence as the force underlying any potential difference among these entities. These results coincide well with an earlier caveat provided by Warren *et al.* (2014), that SDM non-equivalency is not always indicative of ecological speciation, and that is likely over-prescribed in the literature. Regardless, the putative distinctness of *B. berlandieri* warrants further investigation genetically, morphologically, and ecologically.

A greater number of high-resolution, georeferenced, taxonomically-vetted specimen records from across species ranges would allow us to infer higher quality estimates of occupied niche space. Reducing the assumed georeferencing and taxonomic errors in publically available locality datasets would allow use of higher resolution predictor variables, for which many more datasets exist, e.g. Harmonized Soils, which may prove to be important in *Brahea*, given the preference of some species for calcareous soils. These improved models would allow us to include more *Brahea* species, such as *B. calcarea* and *decumbens*, and furthermore to extend these analyses back in time to ancestral nodes.

## Conclusions

We have conducted an explicit phylogenetic study of one of the most poorly understood, yet ecologically important groups of American palms, providing strong resolution and support for phylogenetic relationships in the genus, based on nearly 400 kb of sequence data from all three genomes. We further provide a phylogenetic test of subgeneric species assignments, largely corroborating earlier work based on diagnostic morphological characters. The exception is *B. moorei*, which is strongly supported as being a member of subgenus *Erythea*, but was previously placed in subgenus *Brahea* based on morphological characters including a lack of petiolar spines. We demonstrate the evolution of dwarf forms from taller, tree-like ancestors in *B. moorei* and *B. decumbens.* We provide the first comprehensive estimates of the timing of speciation events across the genus, reveal roles for sympatric speciation and dispersal in the biogeographic history of *Brahea*, and provide evidence of ecological niche divergence among closely related species in the *B. armata* and *B. dulcis* species complexes. The findings are relevant in elucidating complex patterns of biogeographic history in Central America and Mexico. Most importantly, we provide a framework for future studies to be conducted including integrative species delimitation within the genus, estimation of gene flow, phylogeographic history, and more fine-scale investigation of environmental/ecological factors driving evolution in this clade.

## Supplementary Information

Supplementary Link 1 contains MaxEnt outputs of replicated SDMs for *Brahea aculeata*, *B. armata*, *B. berlandieri*, *B. brandegeei*, and *B. dulcis.* Each folder contains an HTML document named ‘maxent.html,’ which details all results generated by MaxEnt for each species. Link: \https://drive.google.com/file/d/1o0Gp-9v7rk2Ou3xVQ7IUelGBaflhZxib/view?usp=sharing.

## Acknowledgements

We thank the staff at the Huntington Library, Art Collections and Botanical Gardens and the University of California Botanical Garden (Holly Forbes) for assistance in collecting material and voucher information. We thank the staff at Global Biologics, LLC. (Sean Blake), Laragen, Inc. (Jinliang Li, Lindy Him), the WVU Genomics Core Facility (Sandy Simon, Ryan Percifield), and the WVU-Marshall Shared Sequencing Facility (Don Primerano, Jun Fan) for sequencing assistance. Financial support was provided by the California State University Program for Education & Research in Biotechnology, and the WVU Program to Stimulate Competitive Research. We thank two anonymous reviewers for their comments improving the manuscript.

